# Use of a human small airway epithelial cell line to study the interactions of *Aspergillus fumigatus* with pulmonary epithelial cells

**DOI:** 10.1101/2023.04.18.537379

**Authors:** Hong Liu, Jianfeng Lin, Quynh T. Phan, Fabrice N. Gravelat, Donald C. Sheppard, Scott G. Filler

**Author notes:** Address correspondence to Hong Liu, or Scott G. Filler,.

## Abstract

During the initiation of invasive aspergillosis, inhaled *Aspergillus fumigatus* conidia are deposited on the epithelial cells lining the bronchi, terminal bronchioles, and alveoli. While the interactions of *A. fumigatus* with bronchial and type II alveolar cell lines have been investigated *in vitro*, little is known about the interactions of this fungus with terminal bronchiolar epithelial cells. We compared the interactions of *A. fumigatus* with the A549 type II alveolar epithelial cell line and the HSAEC1-KT human small airway epithelial (HSAE) cell line. We found that *A. fumigatus* conidia were poorly endocytosed by A549 cells, but avidly endocytosed by HSAE cells. *A. fumigatus* germlings invaded both cell types by induced endocytosis, but not by active penetration. A549 cell endocytosis of *A. fumigatus* was independent of fungal viability, more dependent on host microfilaments than microtubules, and induced by *A. fumigatus* CalA interacting with host cell integrin α5β1. By contrast, HSAE cell endocytosis required fungal viability, was more dependent on microtubules than microfilaments, and did not require CalA or integrin α5β1. HSAE cells were more susceptible than A549 cells to damage caused by direct contact with killed *A. fumigatus* germlings and by secreted fungal products. In response to *A. fumigatus* infection, A549 cells secreted a broader profile of cytokines and chemokines than HSAE cells. Taken together, these results demonstrate that studies of HSAE cells provide complementary data to A549 cells and thus represent a useful model for probing the interactions of *A. fumigatus* with bronchiolar epithelial cells *in vitro*.

**Importance:** During the initiation of invasive aspergillosis, *Aspergillus fumigatus* invades, damages, and stimulates the epithelial cells that line the airways and alveoli. Previous studies of *A. fumigatus*- epithelial cell interactions *in vitro* have used either large airway epithelial cell lines or the A549 type II alveolar epithelial cell line. The interactions of fungi with terminal bronchiolar epithelial cells have not been investigated. Here, we compared the interactions of *A. fumigatus* with A549 cells and the Tert-immortalized human small airway epithelial HSAEC1-KT (HSAE) cell line. We discovered that *A. fumigatus* invades and damages these two cell lines by distinct mechanisms. Also, the proinflammatory responses of the cell lines to *A. fumigatus* are different. These results provide insight into how *A. fumigatus* interacts with different types of epithelial cells during invasive aspergillosis and demonstrate that HSAE cells are useful in vitro model for investigating the interactions of this fungus with bronchiolar epithelial cells.

## Introduction

*Aspergillus fumigatus* is a saprophytic fungus that grows in decaying organic and plant materials (1). It produces airborne spores (asexual conidia) with a diameter of 2-3 µm that are small enough to reach the lower respiratory tract when inhaled (2). In healthy individuals, inhaled conidia are quickly cleared by alveolar microphages and neutrophils (3, 4). In susceptible individuals, inhaled conidia that are not eliminated by these host defenses can adhere to and invade the airways and alveoli, causing invasive aspergillosis (IA). Even with antifungal treatment, this disease remains associated with high morbidity and mortality (5–9). Risk factors for IA include prolonged neutropenia, chemotherapy, corticosteroids, solid organ transplantation, and small molecule inhibitors of myeloid function. Infections with respiratory viruses such as influenza and SARS-CoV-2 that damage the airway epithelium are also a risk factor for IA (10).

Pulmonary epithelial cells are central to the pathogenesis of IA. After *A. fumigatus* conidia are inhaled, they are deposited in the bronchi, terminal bronchioles, and alveoli. In susceptible hosts, *A. fumigatus* conidia progressively form swollen conidia, germlings, and finally mature hyphae, which adhere to, invade and damage host cells (5, 11). *A. fumigatus* conidia and germlings have been found to invade pulmonary epithelial cells *in vitro* by induced endocytosis (11–18). Most studies on the pathogenesis of IA have used the A549 type II alveolar epithelial cell line and the BEAS-2B or 16HBE bronchial cell lines (13, 16, 19–22). Although the small airway epithelial cells that line the terminal bronchioles are likely key targets of *A. fumigatus* during the development of IA, this type of epithelial cell has been studied only in the context of viral infections (23–26). It is important to study bronchiolar epithelial cells because they are transcriptionally and functionally distinct from alveolar and bronchial epithelial cells (27–29).

Previously, we found that integrin α5β1 binds to the *A. fumigatus* CalA invasin and induces fungal endocytosis by A549 cells (16). In the corticosteroid treated mouse model of IA, a Δ*calA* mutant exhibits reduced invasion of the alveolar epithelium but wild-type levels of invasion of bronchiolar and bronchial epithelial cells. These results suggest that *A. fumigatus* interacts differently with alveolar epithelial cells as compared to other pulmonary epithelial cells. To investigate this possibility, we analyzed the interactions of *A. fumigatus* with A549 cells and the Tert-immortalized human small airway epithelial HSAEC1-KT (HSAE) cell line. We discovered that *A. fumigatus* invades and damages these two different cell lines by distinct mechanisms and that the proinflammatory responses of the two cell lines to *A. fumigatus* also differ. These results provide insight into how *A. fumigatus* interacts with different types of pulmonary epithelial cells during the initiation of IA and indicate that HSAE cells are a useful *in vitro* model for investigating the interactions of this fungus with the epithelial cells that line the terminal bronchioles.

## RESULTS

### *A. fumigatus* interacts with both alveolar epithelial cells and small airway epithelial cells during IA in mice

To confirm that *A. fumigatus* interacts with small airway epithelial cells during the initiation of IA, mice were immunosuppressed with cortisone acetate and then intratracheally inoculated with 10^7^ conidia of *A. fumigatus* Af293. After 12 h infection, the lungs were harvested after which thin sections of the lungs were stained with Gomori methenamine silver to visualize the fungi. We observed that the conidia and germinating hyphae were deposited along the entire respiratory tree, including the bronchi, small airways, terminal bronchioles and alveoli (Fig. 1A). These findings suggest that *A. fumigatus* likely interacts with multiple types of pulmonary epithelial cells, including small airway epithelial cells, during the initiation of IA.

**Fig 1.**
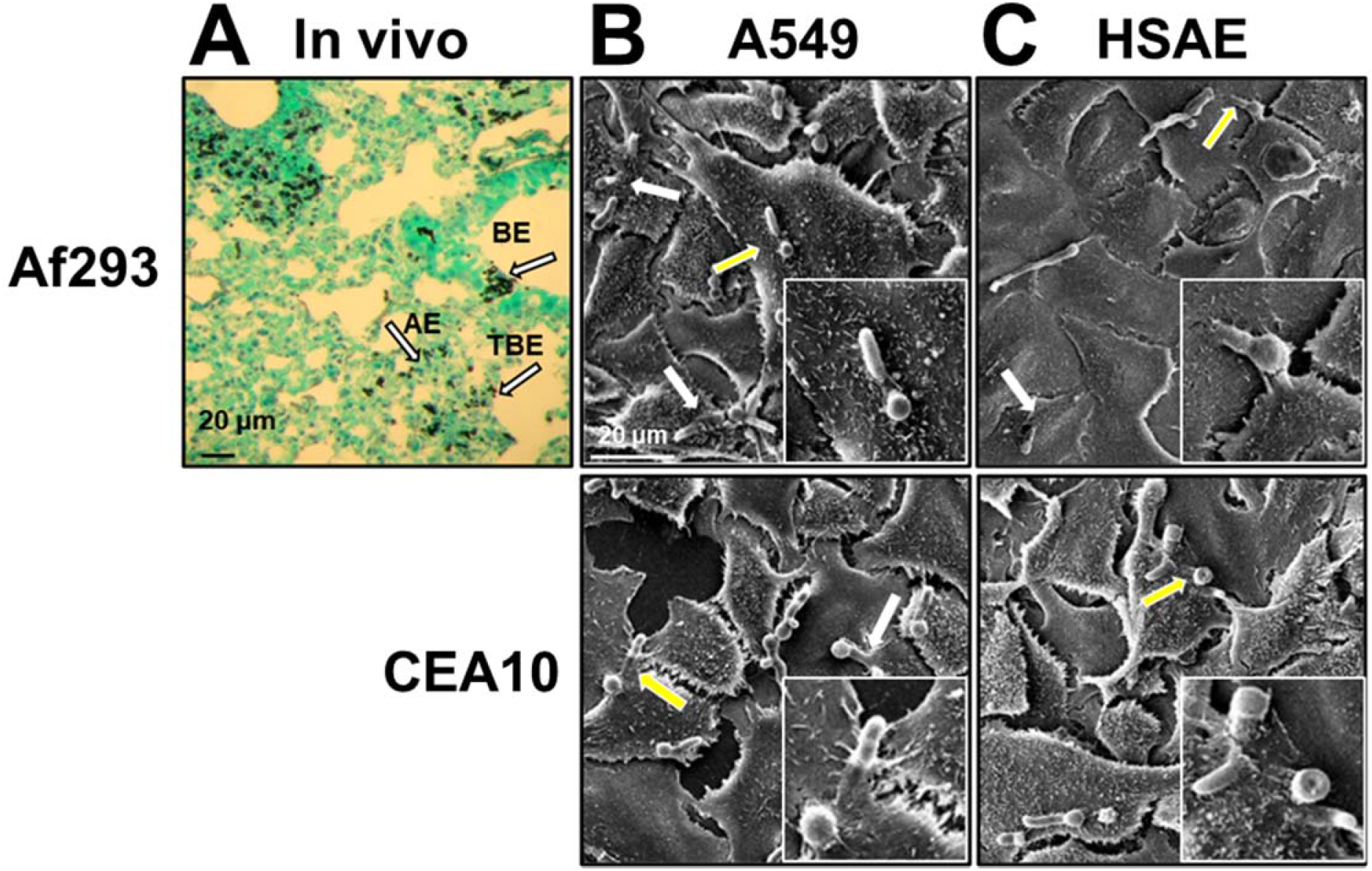
*A. fumigatus* infection in vivo and in vitro. (A) Photomicrograph of Gomori methenamine silver-stained section of a mouse lung after 12-h infection with *A. fumigatus* Af293. Arrows indicate the organisms interacting with bronchial epithelail cells (BE), terminal bronchioloar epithelial cells (TBE), and alveolar epithelial cells (AE). Scale bar 20 µm. (B and C) Scanning electron micrographs of A549 (B) and HSAE (C) cells infected with germlings of *A. fumigatus* strain Af293 (top) or CEA10 (bottom). White arrows indicate the organisms that were endocytosed by A549 or HSAE cells. Yellow arrows indicate the organisms in the magnified images in the lower right panels. Scale bar, 20 µm.

### *A. fumigatus* invades A549 alveolar epithelial cells and HSAE small airway epithelial cells by different mechanisms

To investigate the interactions of *A. fumigatus* with epithelial cells from the alveoli and the terminal bronchioles *in vitro,* we used the A549 type II alveolar epithelial cell line and the HASE human small airway epithelial cell line. We first examined how *A. fumigatus* germlings invade these cell lines using two well characterized *A. fumigatus* clinical isolates, strains Af293 and CEA10. By scanning electron microscopy, we observed that A549 cells preferentially endocytosed the proximal portion of the hyphae, adjacent to the conidia (Fig. 1B). By contrast, HSAE cells endocytosed either the tip of the hyphae or the entire germling, suggesting that the mechanism by which A549 and HSAE cells endocytose *A. fumigatus* may be different (Fig. 1C).

Conidia and germlings are the two major morphotypes of *A. fumigatus* that interact with epithelial cells during early IA development. We used our standard differential fluorescent assay to quantify the endocytosis of *A. fumigatus* conidia and germlings by A549 and HSAE cells. We found that although A549 cells endocytosed few conidia (Fig. 2A), they avidly endocytosed germlings (Fig. 2B). The number of organisms that remained associated with A549 cells after washing, a measure of adherence, was similar for conidia and germlings. HSAE cells responded differently, endocytosing more conidia than germlings (Fig. 2C and D). However, conidia and germlings adhered similarly to these cells. The differences between A549 and HSAE cells were observed with both strains Af293 and CEA10, suggesting that the two types of pulmonary epithelial cells endocytose different *A. fumigatus* morphotypes by different mechanisms.

**Fig 2.**
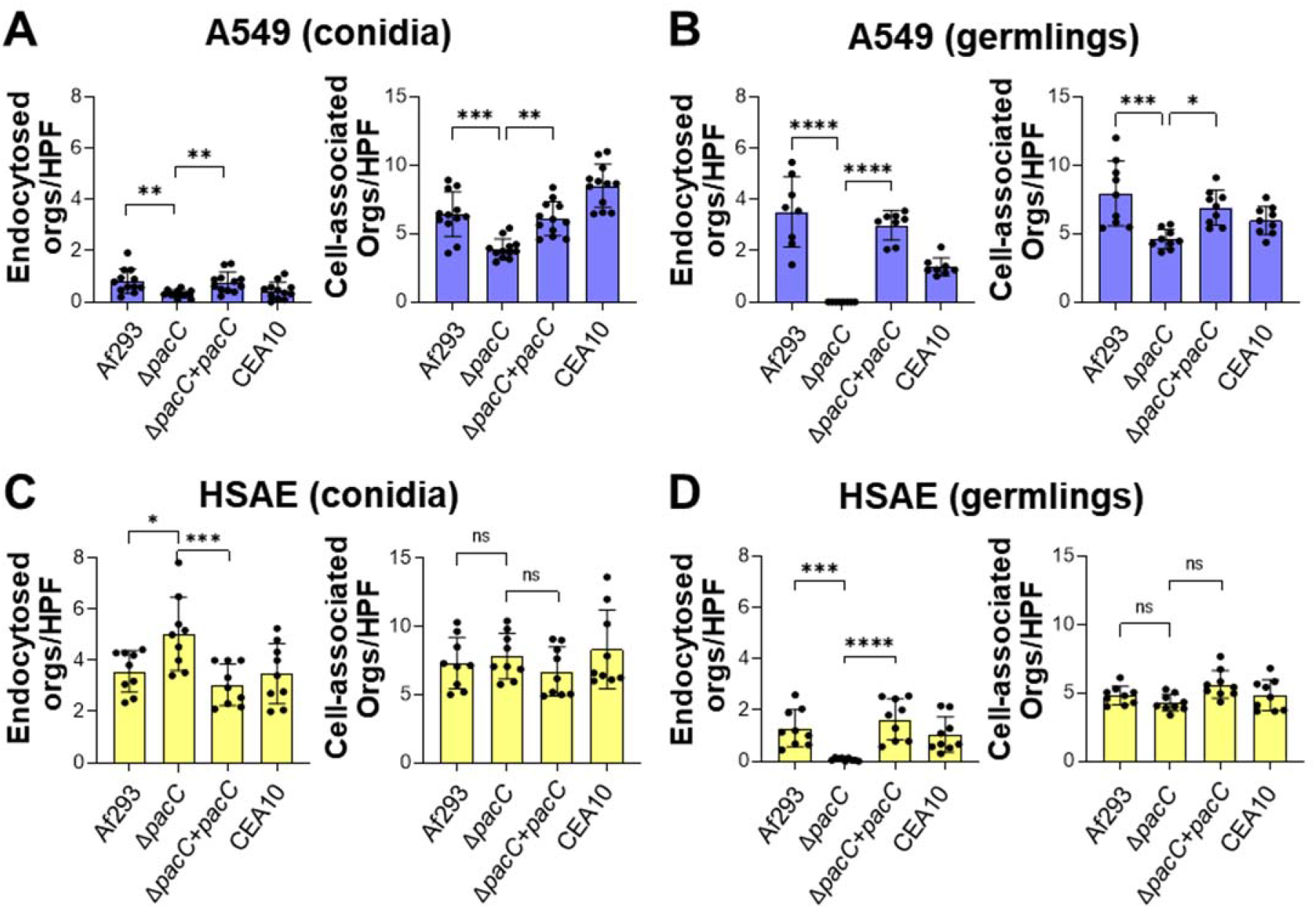
*A. fumigatus* conidia and germlings invades A549 and HSAE cells by different mechanisms. (A and B) A549 cell endocytosis and cell-association (a measure adherence) of conidia (A) and germlings (B) of the indicated *A. fumigatus* strains. (C and D) HSAE cell endocytosis and cell-association of conidia (C) and germlings (D) of the indicated *A. fumigatus* strains. Results are mean ± SD of 3 independent experiments, each performed in triplicate. orgs/HPF, organisms per high-powered field; ns, not significant; \**P* < 0.05; \*\**P* < 0.01; \*\*\**P* < 0.001; \*\*\*\**P* < 0.0001 by ANOVA with the Dunnett’s test for multiple comparisons.

*A. fumigatus* PacC is a pH responsive transcription factor and a Δ*pacC* mutant is defective in invading A549 cells (13). To examine the role of PacC in governing invasion of HASE cells, we constructed a *pacC* gene deletion mutant and corresponding complemented strain in the Af293 strain background. As previously reported, both conidia and germlings of the Δ*pacC* mutant were endocytosed very poorly by A549 cells and this invasion defect was restored with the Δ*pacC+pacC* complemented strain (Fig. 2A and B). The Δ*pacC* mutant also had modestly reduced adherence to A549 cells. Although HSAE cells endocytosed germlings of the Δ*pacC* mutant poorly, they endocytosed Δ*pacC* conidia better than wild-type conidia (Fig. 2C and D). Also, deletion of *pacC* had no effect on adherence to HSAE cells. These results indicate that although PacC is a positive regulator of conidial surface proteins that induce endocytosis by A549 cells, it is a negative regulator of conidial surface proteins that induce endocytosis by HSAE cells. Also, PacC positively regulates of expression of *A. fumigatus* surface proteins that mediate the endocytosis of germlings by both types of epithelial cells.

### *A. fumigatus* invades A549 and HSAE cells by induced endocytosis

Some fungi such as *Candida albicans* can invade host cells by both induced endocytosis and active penetration, in which progressively elongating hyphae physically push their way into host cells (30). To determine if active penetration occurs during *A. fumigatus* invasion, we performed the invasion assay using A549 and HSAE cells that had been fixed with paraformaldehyde prior to infection thereby preventing induced endocytosis. We found that there was no *A. fumigatus* invasion of killed A549 cells or HASE cells by Af293 or CEA10 germlings (Fig. 3A and B). Killing A549 cells resulted in a modest reduction in *A. fumigatus* adherence, but killing HSAE cells had no effect on this interaction. These data indicate that *A. fumigatus* germlings invade alveolar epithelial cells and small airway epithelial cells only by induced endocytosis and not active penetration. Thus, fungal invasin(s) and host cell receptor(s) are crucial for *A. fumigatus* to induce its own endocytosis by both types of pulmonary epithelial cells.

**Fig 3.**
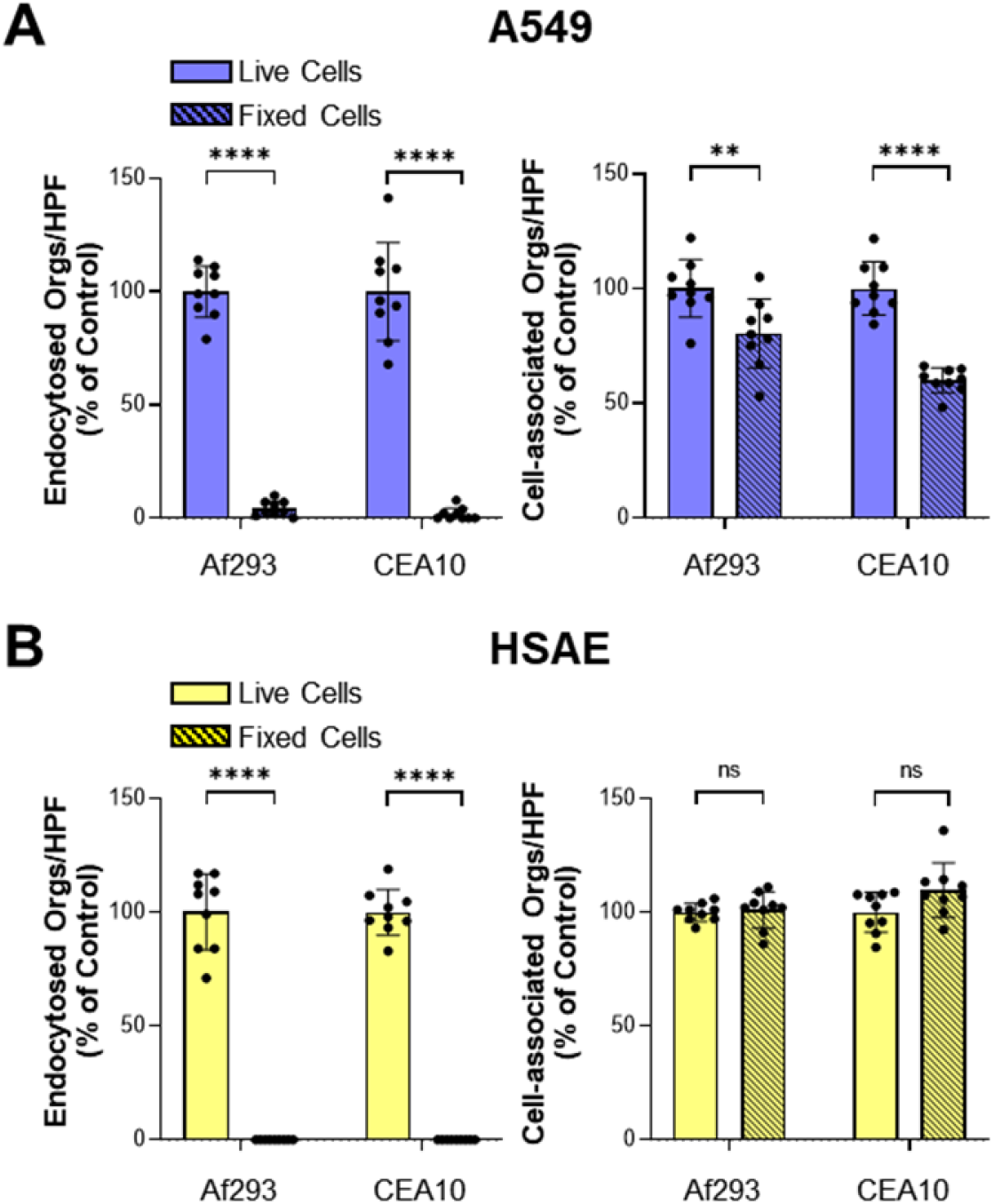
*A. fumigatus* invades both A549 and HSAEC cells by induced endocytosis. (A and B) Endocytosis and cell-association of *A. fumigatus* Af293 and CEA10 by live and paraformaldehyde fixed A549 cells (A) and HSAE cells (B). Results are mean ± SD of 3 independent experiments, each performed in triplicate. orgs/HPF, organisms per high-powered field; ns, not significant; \*\**P* < 0.01; \*\*\*\**P* < 0.0001 by unpaired Students t-test.

### Microfilaments and microtubules play different roles in the endocytosis of *A. fumigatus* by A549 cells vs. HSAE cells

Previously, we found that *A. fumigatus* is endocytosed by A549 cells via an actin-dependent mechanism and that inhibiting polymerization of actin microfilaments with cytochalasin D reduces fungal endocytosis by A549 cells (16). To compare the role of actin in the endocytosis of *A. fumigatus* by A549 and HSAE cells, we infected the cells with *A. fumigatus* germlings and then stained F-actin with ActinRed 555. We observed that actin filaments accumulated around the invading fungal cells in both A549 and HSAE cells (Fig. 4A and 4B). However, there appeared to be more prominent actin accumulation around *A. fumigatus* in A549 cells relative to HSAE cells. This pattern was observed with both *A. fumigatus* Af293 and CEA10.

**Fig 4.**
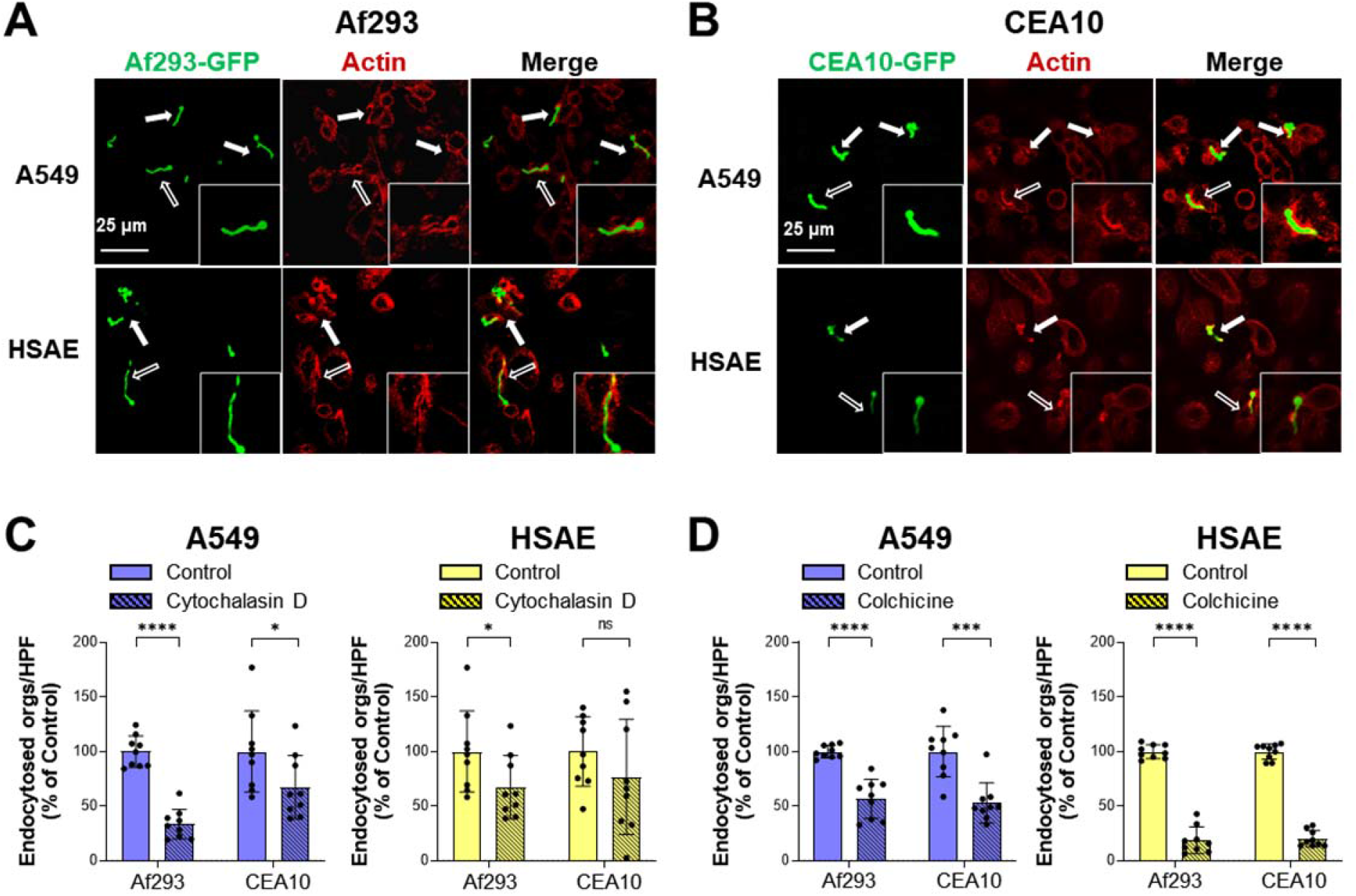
Actin and microtubules play different roles in the endocytosis of *A. fumigatus* by A549 and HSAE cells. (A and B) Confocal images of A549 cells and HSAE cells infected with GFP-expressing strains of Af293 (A) and CEA10 (B) and stained for F-actin (red). Scale bar 24 µm. (C) Effects of 0.6 µM cytochalasin D on the endocytosis of *A. fumigatus* Af293 and CEA10 by A549 and HSAE cells. (D) Effects of 0.5 µM colchicine on the endocytosis of *A. fumigatus* Af293 and CEA10 by A549 and HSAE cells. Results in (C and D) are mean ± SD of 3 independent experiments, each performed in triplicate. orgs/HPF, organisms per high-powered field; ns, not significant; \**P* < 0.05; \*\*\**P* < 0.001; \*\*\*\**P* < 0.0001 by unpaired Students t-test.

To determine the functional significance of the actin accumulation, we investigated the effects of treating A549 and HSAE cells with cytochalasin D. Consistent with our previous results (16), treatment of A549 cells cytochalasin D reduced the endocytosis of strain Af293 by 67% and the endocytosis of strain CEA10 by 33% (Fig. 4C). It had less effect on HSAE cells, reducing the endocytosis of strain Af293 by only 33% and not significantly decreasing the endocytosis of strain CEA10. We also tested the effects of the microtubule polymerization inhibitor, colchicine on the endocytosis of *A. fumigatus*. We found that while treatment of A549 cells with colchicine inhibited endocytosis of strains Af293 and CEA10 by about 40%, it reduced the endocytosis of both strains by HSAE cells by approximately 80% (Fig. 4D). Neither cytochalasin D nor colchicine altered the adherence of *A. fumigatus* to either cell line (Fig. S1) Collectively, these findings indicate that *A. fumigatus* endocytosis by A549 cells is largely dependent on actin microfilaments whereas endocytosis by HASE cells is mainly dependent on microtubules.

### Killed *A. fumigatus* germlings are endocytosed differently by A549 and HSAE cells

Next, we investigated whether fungal viability is required for *A. fumigatus* to adhere to and be endocytosed by A549 and HSAE cells. *A. fumigatus* germlings were killed with the metabolic poison, thimerosal to avoid modifying cell surface ligands. Although killing Af293 germlings decreased their endocytosis by A549 cells by 30%, killing CEA10 germling had no significant effect on their endocytosis (Fig. 5A). Killed organism of both strains had no effect of adherence to A549 cells. By contrast, killing germlings of either strain resulted in an 87-89% reduction in endocytosis by HSAE cells, and slightly increased their adherence (Fig. 5B). Thus, endocytosis of *A. fumigatus* by A549 cells is largely independent of fungal viability, whereas endocytosis by HSAE cells is highly dependent on a factor(s) associated with fungal viability.

**Fig. 5.**
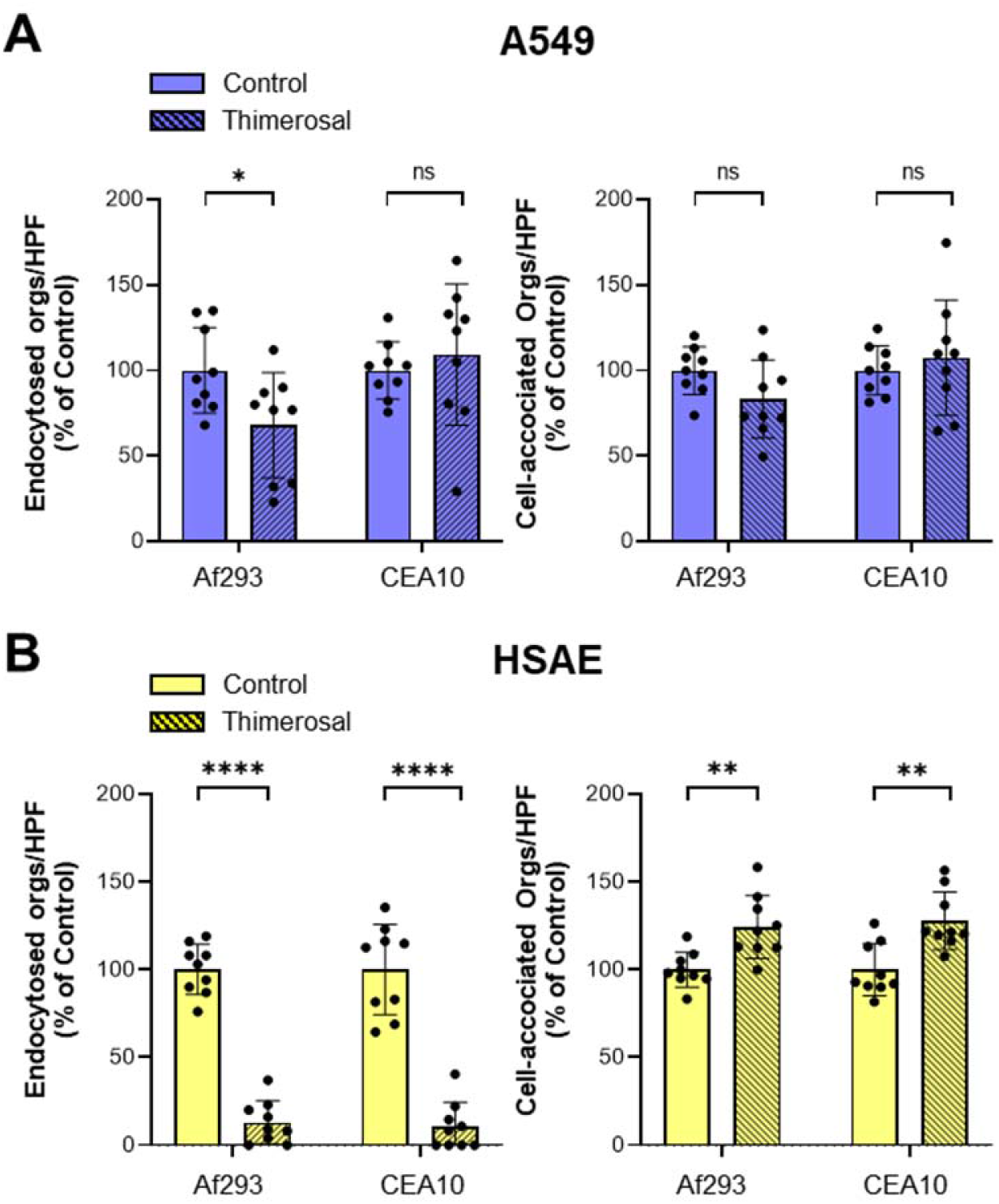
Killed *A. fumigatus* germlings are efficiently endocytosed by A549 cells but not by HSAE cells. Endocytosis and cell-association of live and thimerosal killed germlings of the indicated *A. fumigatus* strains by (A) A549 cells and (B) HSAE cells. Results are mean ± SD of 3 independent experiments, each performed in triplicate. orgs/HPF, organisms per high-powered field; ns, not significant; \**P* < 0.05; \*\**P* < 0.01; \*\*\*\**P* < 0.0001 by unpaired Students t-test.

### *A. fumigatus* endocytosis by A549 cells and HSAE cells is mediated by different fungal invasins and host receptors

Our previous studies with A549 cells showed that the endocytosis of *A. fumigatus* is mediated in part by the interaction of the fungal CalA invasin with the host cell receptor, integrin α5β1 (16). We investigated whether *A. fumigatus* induces its own endocytosis by HSAE cells via the same mechanism. Although the Δ*calA* mutant had significantly reduced endocytosis by A549 cells, it was endocytosed by HSAE cells to levels similar to the wild-type strain (Fig. 6A). The Δ*calA* mutant exhibited wild-type adherence to both cell lines (Fig. S2A). As we have shown previously, blocking integrin β1 and integrin α5 with specific monoclonal antibodies reduced the endocytosis of *A. fumigatus* in A549 cells, however this was not observed with HSAE cells (Fig. 6B). Neither of these antibodies inhibited the adherence of *A. fumigatus* to either cell line (Fig. S2B). To verify this result, we constructed A549 and HSAE cell lines in which *ITGA5* was deleted and in which integrin α5 was no longer expressed (Fig. 6C). Consistent with the antibody inhibition data, deletion of *ITGA5* decreased the endocytosis of *A. fumigatus* by A549 cells but not by HSAE cells and had no effect on adherence to either cell line (Fig. 6D and S2C). Thus, the interactions of *A. fumigatus* CalA with integrin α5β1 induces endocytosis by A549 cells, but not HSAE cells.

**Fig 6.**
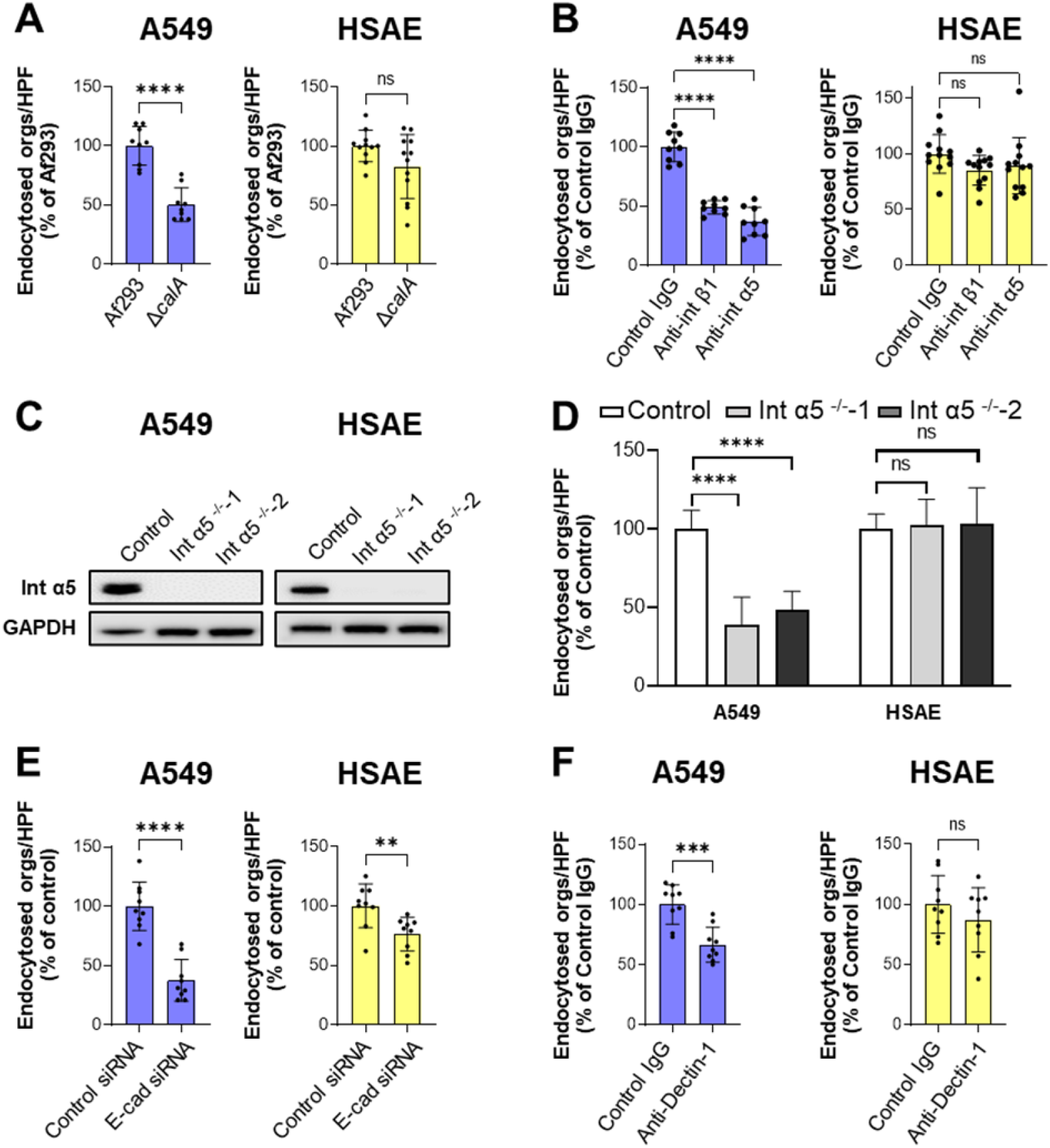
Function of *A. fumigatus* CalA and the host cell receptors α5β1 integrin, E-cadherin and Dectin-1 in mediating endocytosis by A549 and HSAE cells. (A) Endocytosis of wild-type and Δ*calA* strains of *A. fumigatus* by A549 and HSAE cells. (B) Effects of anti-β1 integrin and anti-α5 integrin antibodies on the endocytosis of *A. fumigatus* Af293 by A549 and HSAE cells. (C) Western blots showing the deletion of integrin α5 (Int α5) in two different clones of A549 and HSAE cells. (D) Deletion of integrin α5 inhibits the endocytosis of *A. fumigatus* by A549 cells but not HSAE cells. (E) Effects of siRNA knockdown of E-cadherin on the endocytosis *A. fumigatus* Af293 by A549 and HSAE cells. (F) Effects of an anti-Dectin-1 antibody on the endocytosis of *A. fumigatus* Af293 by A549 and HSAE cells. Results in (A, B, D, E and F) are the mean ± SD of 3 independent experiments, each performed in triplicate; orgs/HPF, organisms per high-powered field; ns, not significant; ** *P* < 0.01, \*\*\**P* < 0.001, \*\*\*\**P* < 0.0001 by the unpaired Students t-test (A, E and F) and ANOVA with Dunnett’s test for multiple comparison (B and D).

Additional receptors have been reported to mediate the endocytosis of *A. fumigatus* germlings by A549 cells including E-cadherin and Dectin-1 (13, 18). We investigated whether these receptors are also required for HSAE cells to endocytose *A. fumigatus* germlings. Knockdown of E-cadherin with siRNA significantly reduced the endocytosis of *A. fumigatus* by both types of epithelial cells and reduced adherence to A549 cells but not HSAE cells (Fig. 6 E and S2D). By contrast, a neutralizing anti-Dectin-1 antibody significantly inhibited the endocytosis of *A. fumigatus* by A549 cells but had no detectable effect on endocytosis by HSAE cells (Fig. 6F). This antibody did not alter adherence to either cell type (Fig. S2E). Collectively, these results indicate that although E-cadherin is required for the maximal endocytosis of *A. fumigatus* by both A549 cells and HSAE cells, Dectin-1 is only required for endocytosis by A549 cells. The finding that siRNA knockdown of E-cadherin resulted in only a modest decrease in endocytosis by HSAE cells suggests that additional, as yet undefined receptors must mediate the endocytosis of *A. fumigatus* by HSAE cells.

### *A. fumigatus* damages A549 and HSAE cells by distinct mechanisms

*A. fumigatus* causes significant damage to host cells and we have found that the capacity of *A. fumigatus* mutants to damage A549 cells *in vitro* directly correlates with their virulence in the immunosuppressed mouse model of IA (13, 16, 31, 32). We compared the capacity of *A. fumigatus* to damage A549 cells and HSAE cells using our standard ^51^Cr release assay (13, 32). Using strains Af293 and CEA10, we found that live *A. fumigatus* caused less damage to A549 cells relative to HSAE cells (Fig. 7A and B). When the epithelial cells were incubated with germlings that had been killed with either thimerosal or paraformaldehyde, the extent of damage to both cell types was significantly reduced. However, killed germlings, especially those of strain Af293 caused significantly more damage to HSAE cells than to A549 cells. To investigate the role of soluble factor in *A. fumigatus* with the epithelial cells, we incubated them with filtrates from cultures of *A. fumigatus* cells grown in tissue culture medium for 48 h. Although filtrates prepared from cultures of strain Af293 caused more damage to HSAE cells than A549 cells the first 6 h of exposure, these filtrates killed all of the HSAE and A549 cells by 24 h (Fig. 7C). Culture filtrates prepared from strain CEA10 caused no detectable damage either HSAE or A549 cells for the first 6 h of incubation. However, they caused a modest amount of damage to HSAE cells and no detectable damage to A549 cells after 24 h (Fig. 7D). Collectively, these results suggest that HSAE cells are more susceptible than A549 cells to damage caused by direct contact with both live and killed *A. fumigatus* cells and by products that are secreted by this fungus. These data also indicate that under the conditions used for these assays, although live cells of strain Af293 damaged epithelial cells similarly to strain CEA10, killed germlings and culture filtrates of strain Af239 caused greater epithelial cell damage than the other strain.

**Fig. 7.**
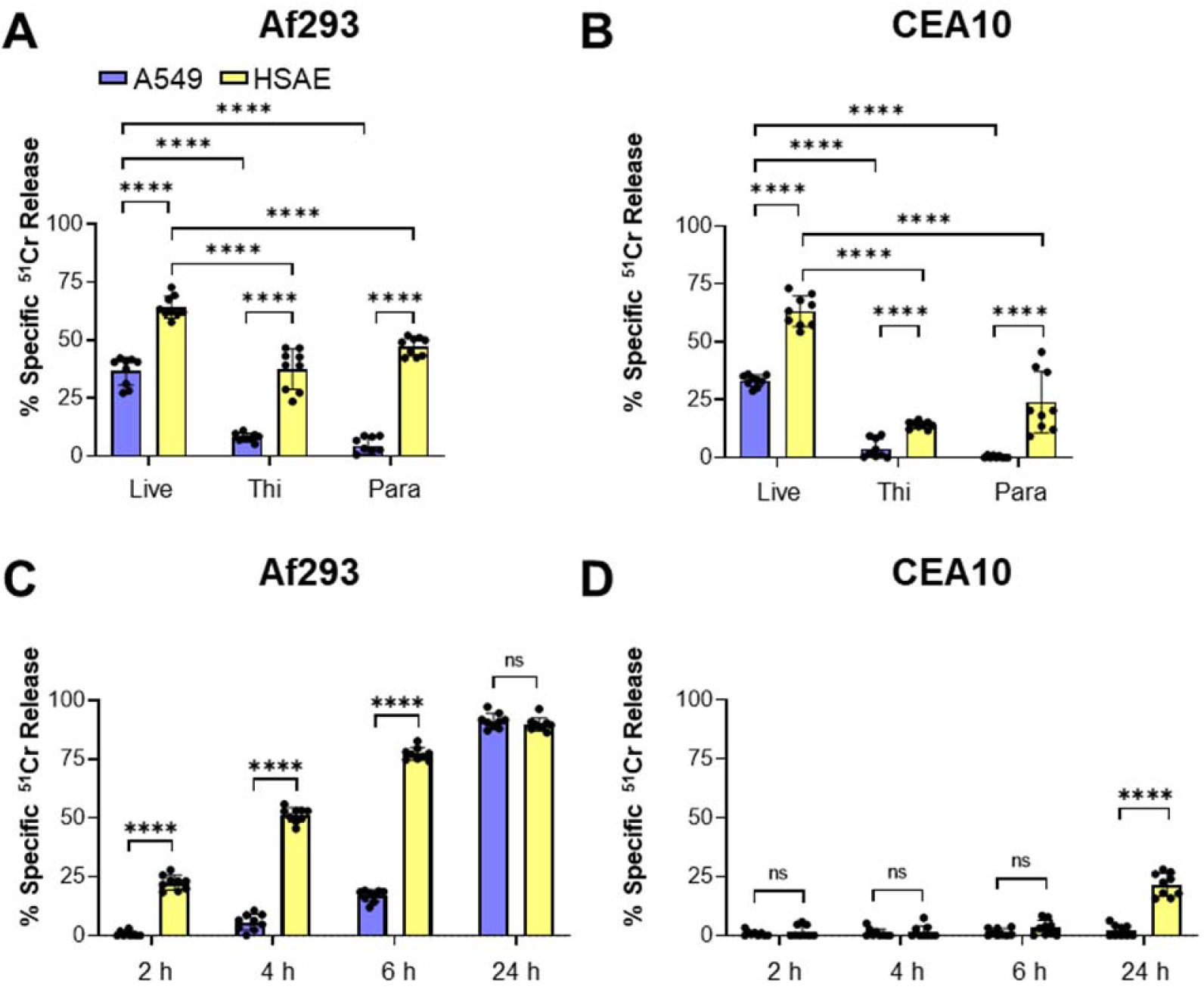
Fungal viability and secreted products have different effects on *A. fumigatus*- induced damage of A549 and HSAE cells. (A and B) Damage to A549 and HSAE cells caused by *A. fumigatus* strains Af293 (A) or CEA10 (B). Damage was measured after a 24 h exposure to live germlings or germlings that had been killed with either thimerosal (Thi) or paraformaldehyde (Para). (C and D) Time course of damage to A549 and HSAE cells by culture filtrates of strains Af293 (C) or CEA10 (D). *A. fumigatus* cells were grown in tissue culture medium for 48 h at 37°C after which the conditioned medium was filter sterilized, mixed with fresh medium at a ratio of 1:3, and added to A549 and HSAE cells. Results are mean ± SD of 3 independent experiments, each performed in triplicate; ns, not significant; \*\*\*\**P* < 0.0001 by ANOVA with the Dunnett’s test for multiple comparisons.

### *A. fumigatus* infection stimulates different pro-inflammatory responses in A549 and HSAE cells

Pulmonary epithelial cells play a central role in orchestrating the host defense against pulmonary pathogens by secreting pro-inflammatory cytokines and chemokines that attract and activate professional phagocytes that kill the invading microorganism (33–36). To compare the inflammatory responses of A549 and HSAE cells to *A. fumigatus*, we infected them for 24 h with conidia of two different strains of *A. fumigatus* and measured the levels of 8 cytokines and chemokines using a Luminex Multiplex Array. We observed different patterns of response that varied with the type of pulmonary epithelial cell. *A. fumigatus* infection stimulated both A549 cells and HSAE cells to secrete increased amounts of CXCL8 and IL-6 relative to uninfected cells, but A549 cells secreted greater amounts of these pro-inflammatory mediators than HSAE cells (Fig. 8). While *A. fumigatus* stimulated A549 cells to secrete TNF-α, GM-CSF, and CCL2, the secretion of these mediators by HSAE cells was undetectable in both infected and uninfected cells. *A. fumigatus* stimulated both A549 cells and HSAE cells to secrete similar amounts of IL-1α and IL-1β. CXCL1 secretion was induced by infection of HSAE cells, but not A549 cells. Collectively, these results suggest that while both cell types can potentially influence the host immune response to *A. fumigatus*, A549 cells produce a greater repertoire of cytokines and chemokines than HSAE cells.

**Fig 8.**
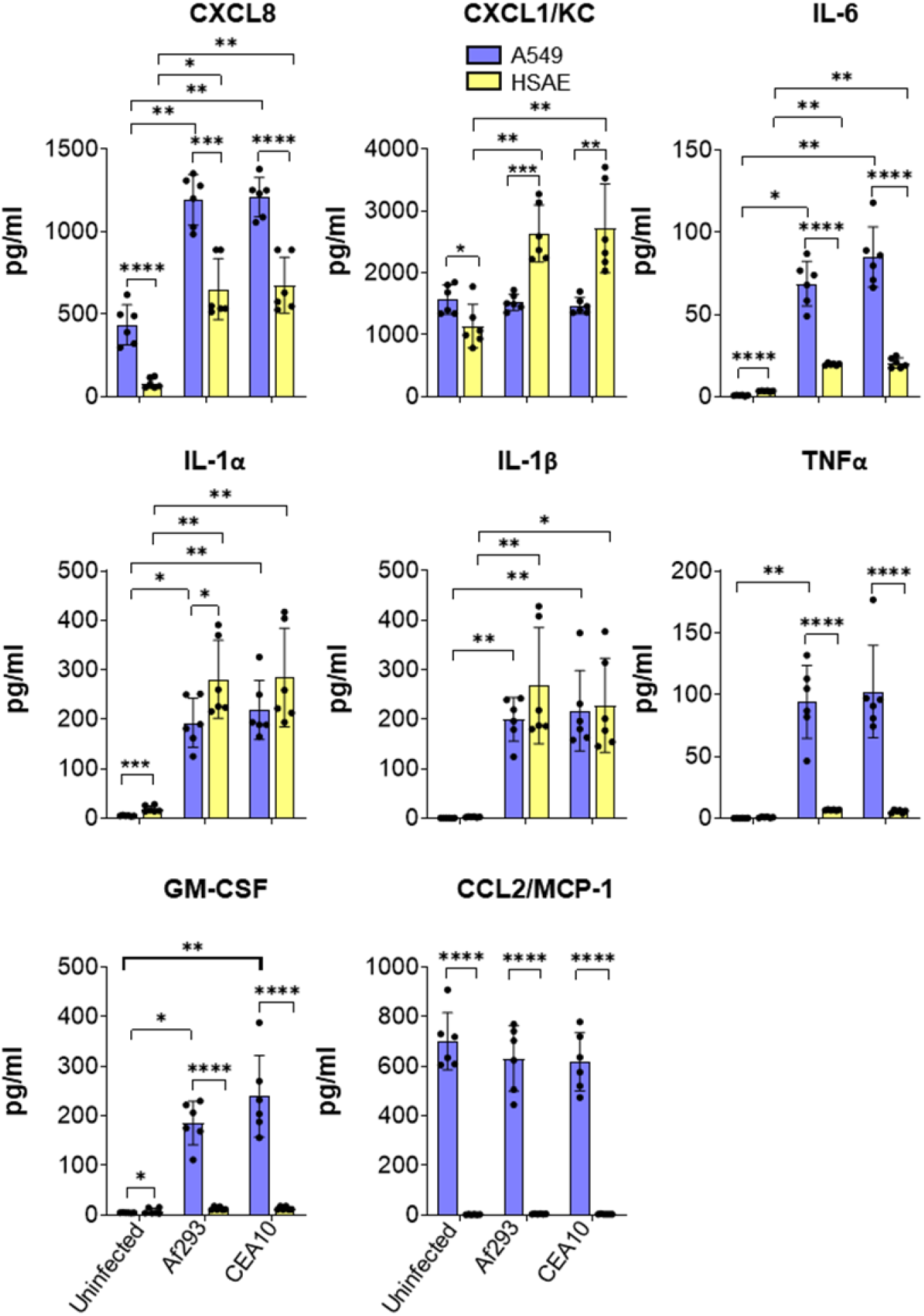
Different pro-inflammatory responses of A549 and HSAE cells to *A. fumigatus* infection. A549 and HSAE cells were infected with Af293 or CEA10 strains for 24 h, after which the conditioned medium was collected, and the levels of the indicated cytokines and chemokine were measure by Luminex Multiplex Array. Results are mean ± SD of 3 independent experiments, each performed in duplicate. \**P* < 0.05, \*\**P* < 0.01, \*\*\**P* < 0.001, \*\*\*\**P* < 0.0001 by ANOVA with the Dunnett’s test for multiple comparisons.

## Discussion

During the initiation of IA, inhaled *A. fumigatus* conidia likely interact with the pulmonary epithelial cells that line the entire respiratory tree, including terminal bronchiolar epithelial cells and alveolar epithelial cells. In this study, we determined that *A. fumigatus* interacts differently with these two types of pulmonary epithelial cells *in vitro* (Summarized in Table S1). One major difference was the degree of endocytosis of conidia by the two different cell lines. While A549 cells endocytosed conidia poorly, HASE cells avidly endocytosed this morphotype. Also, fewer Δ*pacC* conidia were endocytosed by A549 cells relative to wild-type conidia, whereas more Δ*pacC* conidia than wild-type conidia were endocytosed by HSAE cells. These results suggest that different receptor-ligand interactions induce conidial endocytosis by A549 cells as compared to HSAE cells.

Previously, it has been reported that A549 cells avidly endocytosis *A. fumigatus* conidia, in contrast to the results presented here (13, 22, 37). A key difference is that the previous endocytosis assays were performed in the presence of serum, whereas the current ones were performed in serum-free media to better mimic the conditions to which the fungus is exposed *in vivo.* Nevertheless, the finding that serum enhances the endocytosis of *A. fumigatus* suggests that serum proteins may function as bridging molecules between conidia and alveolar epithelial cells, and thereby induce endocytosis. Recently, we found that high molecular weight kininogen and vitronectin in human serum can act as bridging molecules between yeast-phase *Candida* spp. and vascular endothelial cells (38). Whether these proteins can also function as bridging molecules between *A. fumigatus* conidia and alveolar epithelial cells is currently unknown.

*C. albicans* can invade oral epithelial cells by two distinct mechanisms, active penetration and induced endocytosis (30, 39, 40). Induced endocytosis occurs when the *C. albicans* invasins, Als3 and Ssa1 interact with multiple oral epithelial cell receptors, including E-cadherin, the epidermal growth factor receptor, HER2, and the ephrin type-A receptor 2 (39–43). This process requires intact microfilaments (44). We determined that *A. fumigatus* germlings invade A549 cells and HASE cells only by induced endocytosis and not by active penetration. However, the mechanisms of induced endocytosis by A549 cells and HSAE cell are different. Endocytosis of *A. fumigatus* by A549 cells is similar to the endocytosis of *C. albicans* by oral epithelial cells in that it is mediated in part by E-cadherin, does not require fungal viability and is highly dependent on host microfilaments. Unlike the endocytosis of *C. albicans,* endocytosis of *A. fumigatus* by A549 cells is induced by Dectin-1 and by CalA interacting with host cell integrin α5β1 (16). By contrast, HSAE cell endocytosis is different from epithelial cell endocytosis of *C. albicans* in that it requires fungal viability, is more dependent on microtubules than microfilaments, and minimally dependent on E-cadherin. HSAE cell endocytosis of *A. fumigatus* also differs from that of A549 cells because it independent of CalA, integrin α5β1, and Dectin-1. The mechanisms by which *A. fumigatus* induces its own endocytosis by HSAE cells is currently unknown but is topic of active investigation.

The capacity of *A. fumigatus* to damage host cells is a key aspect of the pathogenesis of IA. Thus, it was of interest to determine that HSAE cells were more susceptible than A549 cells to damage caused by live and killed *A. fumigatus* germlings and by *A. fumigatus* culture filtrates. We also found that killed germlings and culture filtrates of strain Af293 caused more damage to both types of epithelial cells relative to strain CEA10. While the exact mechanisms by which *A. fumigatus* damages host cells are incompletely understood, it is virtually certain that damage is caused by secondary metabolites produced by the fungus. In *Aspergillus spp.,* the genes that encode the enzymes for secondary metabolite production are usually located in gene clusters that exhibit significant strain-to-strain variability. The genome of strain Af293 contains 35 biosynthetic gene clusters whereas that of strain CEA10 contains 33, and the profile of secondary metabolites produced by strain Af293 *in vitro* is different from that of CEA10 (45). Also, we have found that the SltA transcription factor governs the expression of different biosynthetic gene clusters in strain Af293 relative to strain CEA10 (32). These genetic differences likely account for the differences in epithelial cell damage caused by the two *A. fumigatus* strains.

Pulmonary epithelial cells provide the first-line defense against fungal pathogens during lung infection. In addition to functioning as a physical barrier to infection, they secrete cytokines and chemokines that recruit and activate phagocytes to kill the invading fungus (17, 36, 46). Although we determined that both A549 and HSAE cells responded to *A. fumigatus* infection by secreting proinflammatory cytokines and chemokines, the patterns of these response were different. Overall, A549 cells secreted higher levels and a greater range of these immunomodulators than HSAE cells, suggesting that type II alveolar cells may play a greater role than small airway epithelial cells in orchestrating the host response to *A. fumigatus.* However, because small airway epithelial cells are more numerous than type II alveolar cells, they still may play a significant role in the host defense against *A. fumigatus*. This hypothesis needs to be evaluated by *in vivo* studies.

During the pathogenesis of invasive aspergillosis, the fungus interacts with multiple types of pulmonary epithelial cells. To develop therapeutic strategies to block the capacity of *A. fumigatus* to invade and damage the pulmonary epithelium, it is necessary to develop a comprehensive understanding of how the fungus interacts with the different types of epithelial cells that line the airways and alveoli. Here, we demonstrate that *A. fumigatus* invades and damages A549 and HSAE cells by different mechanisms and that these two different cell lines produce different profiles of pro-inflammatory mediators in response to fungal infection. Thus, HSAE cells provide complementary data to A549 cells and represent a useful model for studying the interactions of *A. fumigatus* with bronchiolar epithelial cells *in vitro*.

## MATERIALS AND METHODS

### Strains, growth condition

All *A. fumigatus* strains (listed in Table S2) were grown on Sabouraud dextrose agar (Difco) at 37 °C for 7-10 days prior to use. Conidia were harvested with PBS containing 0.1% Tween 80 (Sigma-Aldrich), filtered with a 40 µm cell strainer (Corning), and enumerated with a hemacytometer. To produce germlings, the conidia were incubated in Sabouraud dextrose broth (Difco) at 37°C for 5.5 h as described (16). For experiments with killed germlings, conidia were incubated in Sabouraud dextrose broth for 8 h and then fixed with 4% paraformaldehyde at room temperature for 20 min or with 0.2% of thimerosal at 4°C overnight. Killing was confirmed by absence of growth after incubating the hyphae on Sabouraud dextrose agar plates for 2 d.

### Strain construction

To construct the Δ*pacC* (Afu3g11790) deletion mutant, a transient CRISPR-Cas9 gene deletion system was used (47, 48). The Cas9 expression cassette was amplified from plasmid pFC331, using primers Cas9-F and Cas9-R. All primers are listed in Table S3. To construct the sgRNA expression cassette, two DNA fragments were amplified from plasmid pFC334 using primers sgRNA-F and sgRNA-PacC-R, and sgRNA-R, sgRNA-PacC-F. Next, the sgRNA expression cassette was amplified by fusion PCR from the two DNA fragments, using primers sgRNA-F and sgRNA-R. The hygromycin resistance (HygR) repair template was amplified from plasmid pVG2.2-hph using primers Hyg-F and Hyg-R, which had about 50 bp of homology to the 5’ end of the protein coding sequence of the gene and the 3’ end of the protein coding sequence, respectively. The HygR repair template was mixed with the Cas9 cassette and the two sgRNA cassettes and then used for protoplast transformation. Hygromycin resistant clones were screened for deletion of *pacC* by colony PCR using primers PacC-Screen-F, PacC- Screen-R and insertion of HygR using primers PacC-Screen-up, Hyg-Screen-R. The positive clones were also confirmed for absence of integration of DNA encoding Cas9 or the gRNA, using primers Cas9-ScreenF and Cas9-ScreenR, and sgRNA-ScreenF, sgRNA-ScreenR.

To construct the Δ*pacC* + *pacC* complemented strain, a 5888 bp fragment containing the *pacC* protein coding sequence and approximately 2.5 kb of 5’ flanking sequence and 0.5 kb of 3’ flanking sequence was PCR-amplified from Af293 genomic DNA using primers PacC-Com-F and PacC-Com-R. The resulting fragment was cloned into the NotI site of plasmid p402 (31), which was used to transform the Δ*pacC* strain. To confirm the presence of the complementation plasmid, the phleomycin-resistant colonies were screened by colony PCR using primers PacC- Com-F and PacC-Com-R to detect *pacC*. The transcript levels of *pacC* in the various clones were quantified by real-time RT-PCR using primers PacC-Realtime-F and PacC-Realtime-R. The clones in which the transcript level of *pacC* was most similar to that of the wild-type strain was used in all experiments.

Different strains of *A. fumigatus* that constitutively expressed GFP were constructed to use in epithelial cell invasion assays (32). The Δ*pacC* mutant was transformed with plasmid GFP- Phleo and the Δ*pacC+pacC* complemented strain was transformed with plasmid GFP-pPTRI as previously described (16).

### Cell lines

All cell lines were purchased from a from the American Type Culture Collection. The A549 cell line was cultured in F-12 K medium (30-2004; ATCC) supplemented with 10% fetal bovine serum (FBS) (Gemini Bio-Products) and 1% streptomycin and penicillin (Irvine Scientific). HSAEC1-KT cells were cultured in SAGM BulletKit medium (CC-3119 and CC-4124; Lonza) and HEK293T cells were cultured in DMEM medium (10569010; Gibco) supplemented with 10% heat inactivated FBS and 1% streptomycin and penicillin. Cells were cultured at 37°C with 5% CO_2_. The A549, HSAEC1-KT and HEK293T cell lines were authenticated by ATCC. All cell lines had no mycoplasma contamination. All the experiments in this study were performed using monolayers of A549 and HSAE cells exposed to *A. fumigatus* in serum-free medium. Where relevant, A549 or HSAE cells were fixed with 4% paraformaldehyde (Sigma-Aldrich) for 20 min and then rinsed extensively with PBS before use.

### Ethics statement

The mouse study and experimental procedures were approved by the Animal Care and Use Committee at the Lundquist Institute for Biomedical Innovation at Harbor-UCLA Medical Center.

### Mouse models of invasive aspergillosis

To visualize the epithelial cell interactions of inhaled conidia in vivo, a minor modification of our standard mouse model of invasive aspergillosis was used (16). Briefly, 6-week male Balb/C mice (Taconic Laboratories) were immunosuppressed with cortisone acetate (Sigma-Aldrich; 500 mg/kg) administrated subcutaneously every other day starting at day -4 before infection. Three mice were intratracheally inoculated with 10^7^ conidia of strain Af293. After 12 h of infection, the mice were sacrificed, after which their lungs were harvested and processed for histochemical analysis with Gomori methenamine silver staining. The slides were viewed by light microscopy and representative images were obtained.

### Scanning electron microscopy

For electron microscopy, 10^5^ germlings/ml of strains Af293 and CEA10 were incubated with A549 and HSAE cells on glass coverslips for 2.5 h. The coverslips were then rinsed in PBS and fixed overnight with 2.5% glutaraldehyde in 0.1 M sodium cacodylate buffer at 4°C. After dehydrating and critical-point drying the samples, they were sputter coated with Au-Pd and imaged with a Hitachi S-3000 N scanning electron microscope.

### Host cell endocytosis assay

The endocytosis of conidia and germlings of different strains by A549 and HSAE cell lines was determined by our previously described differential fluorescence assay (16, 32). Briefly, 2.5 x 10^5^ host cells were grown to confluency on glass coverslips in a 24-well tissue culture plate. The cells were incubated with 10^5^ cells of GFP-expressing conidia or germlings and incubated for 6 h for experiments with conidia, 2.5 h for experiments with germlings and live host cells, and 3.5 h for experiments with germlings and killed host cells. At the end of the incubation period, the cells were rinsed with PBS in a standardized manner, fixed with 4% paraformaldehyde, and stained with a rabbit polyclonal anti- *Aspergillus* antibody (Meridian Life Science, Inc.) followed by an AlexaFluor 568-labeled goat anti-rabbit antibody (Life Technologies). The coverslips were mounted inverted on microscope slides and viewed under epifluorescence. At least 100 organisms per coverslip were scored for endocytosis and each strain was tested in triplicate in three independent experiments.

In some assays, the epithelial cells were incubated with 0.6 µM cytochalasin D (C2618; Sigma-Aldrich) or 0.5 µM colchicine (PHR1764; Sigma-Aldrich) in 0.1% DMSO. Control cells were incubated with 0.1% DMSO only. For antibody blocking experiments, the epithelial cells were incubated with 10 µg/mL of an anti-β1 integrin antibody (6S6; Millipore), 25 µg/mL of an anti-α5 integrin antibody (NKI-SAM-1; Millipore), or 3 ug/ml of an anti-Dectin-1 antibody (MAB1859; R&D Systems). Mouse IgG was used as a control. In all experiments, the inhibitors or antibodies were added 45 min prior to infection and remained in the medium for the duration of the experiment.

To knockdown E-cadherin, A549 and HSAE cells were transfected with E-cadherin siRNA (sc-35242; Santa Cruz Biotechnology), or random control siRNA (Qiagen) using Lipofectamine 2000 (Invitrogen) following the manufacturer’s instructions. The efficiency of siRNA knockdown was verified by immunoblotting of whole cell lysates with an anti-E-cadherin (24E10, Cell signaling) antibody.

### CRISPR/Cas9 deletion of integrin α5 in A549 and HSAE cells

Two independent guide RNAs for *ITGA5* (Table S2) from the Brunello sgRNA library (49) were synthesized (Sigma-Aldrich) and cloned in to the pLentiCRISPRv2-blast plasmid (#98293; Addgene). After confirmation by Sanger sequencing, each plasmid was used the generate a lentivirus. Lentiviral packaging was carried out in HEK293T cells using the 3^rd^ generation packaging plasmids pCMV delat R8.2 (#12263; Addgene) and pCMV-VSVG (#8454; Addgene). A549 or HSAE cells were transduced with the lentiviruses in the presence of 10 µg/mL polybrene (Santa Cruz Biotechnology). The next day, transduced cells were selected with 20 µg/ml blasticidin (A11139- 03; Gibco) and the blasticidin-resistant cells were processed for single colonies. Independent single-cell derived gene-deletion cell lines were confirmed by Western blotting with an anti-integrin α5 antibody (AB150361; Abcam).

### Confocal microscopy

To visualize actin remodeling during *A. fumigatus* endocytosis, A549 and HSAE cells were infected for 2.5 h and then fixed as in the endocytosis assay. After permeablizing the cells with 0.1% triton X-100, they were sequentially stained with the anti-aspergillus antibody, AlexaFluor 647-labeled goat anti-rabbit antibody, and ActinRed^TM^ 555 (R37112; Invitrogen). The coverslips were mounted onto slides and imaged by confocal microscopy (Leica SP8).

### Cell damage assay

The amount of damage to A549 and HSAE cells caused by direct contact with *A. fumigatus* was determined using a ^51^Cr release assay as described previously (13). The inoculum was 5 x 10^5^ conidia per well and the incubation time was 24 h. To determine extent of damage caused by killed *A. fumigatus* germlings, the cells were infected with 5 x 10^5^ germlings that had been killed with thimerosal or paraformaldehyde. To determine the amount of damage caused by secreted fungal products, conidia of *A. fumigatus* were incubated in DMEM (Gibco^TM^)at a final concentration of 5 x 10^5^ cells/ml in 5% CO_2_ at 37°C. After 48 h, the culture supernatant was filter sterilized, mixed with fresh tissue culture medium at a ratio of 1:3 (v/v) and added to ^51^Cr-loaded epithelial cells for various times. Each experiment was performed in triplicate at least 3 times.

### Cytokine and chemokine production

A549 and HSAE cells in 24-well plates were infected with 5 x 10^5^ conidia of strain Af293. After 24 h infection, the conditioned medium was collected, clarified by centrifugated, and stored in aliquots at −80 °C. At a later time, the concentration of CXCL8, CXCL1/KC, IL-6, IL-1α, IL-1β, TNFα, GM-CSF, and CCL2/MCP-1 in the medium was determined using the Luminex multiplex assay (R&D Systems). The experiment was performed in duplicate on 3 separate occasions.

### Statistics

All statistical analyses were performed using GraphPad Prism version 8. Data were compared using the unpaired, two-sided Students *t* test or one-way ANOVA with Dunnett’s test for multiple comparisons. *P* values ≤ 0.05 were considered to be statistically significant.

## ACKNOWLEDGEMENTS

This work was supported by NIH grant R01AI162802. We thank Adam Diab for assistance with tissue culture.

---

**Figure S1.**
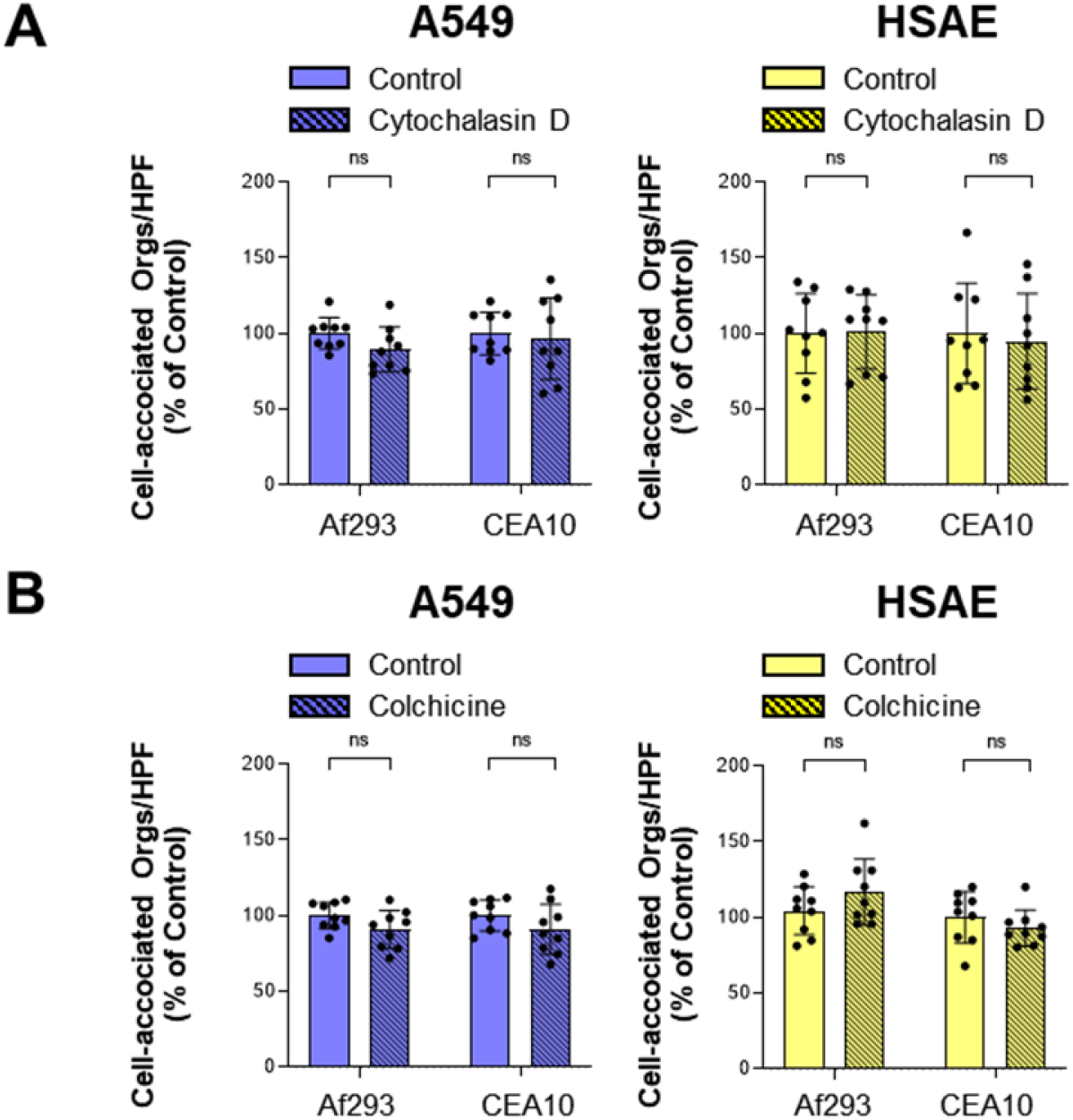
Adherence *A. fumigatus* to A549 and HSAE cells is not affected by cytochalasin D and colchicine. (A) Effects of 0.6 μM Cytochalasin D and (B) 0.5 μM colchicine on the cell-association (a measure of adherence) of the indicated strains of *A. fumigatus* with A549 and HSAE cells. Results are mean ± SD of 3 independent experiments, each performed in triplicate. orgs/HPF, organisms per high-powered field; ns, not significant by the unpaired Students t-test.

**Figure S2.**
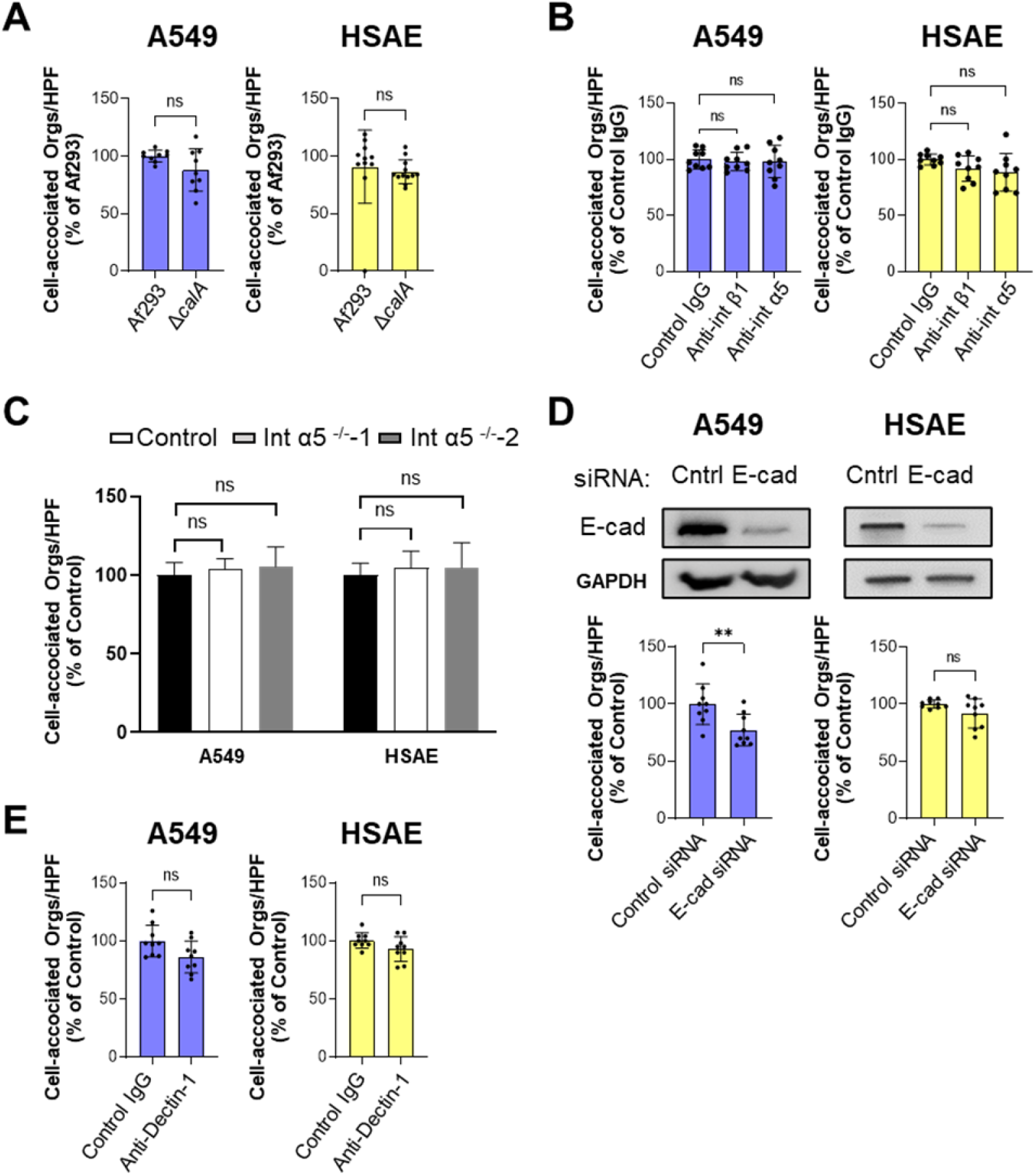
Adherence of *A. fumigatus* to A549 and HSAE cells. (A) Adherence of the indicated *A. fumigatus* strains to A549 and HSAE cells. (B) Effects of anti-β1 integrin and anti-α5 integrin antibodies on the adherence of *A. fumigatus* Af293 to A549 and HSAE cells. (C) Deletion of integrin α5 has no effect on the adherence of *A. fumigatus* Af293 to A549 and HSAE cells. (D) Effects of siRNA knockdown of E-cadherin on the adherence of *A. fumigatus* Af293 to A549 and HSAE cells. Top panels contain representative immunoblots of whole cell lysates showing siRNA knockdown of E-cadherin in A549 and HSAE cells. (E) Effects of anti-Dectin-1 antibody on the adherence of *A. fumigatus* Af293 to A549 and HSAE cells. Results are mean ± SD of 3 independent experiments, each performed in triplicate. orgs/HPF, organisms per high-powered field; ns, not significant; \*\**P* < 0.01 by unpaired Students t-test (A, D and E) and ANOVA with Dunnett’s test for multiple comparison (B and C).

**Table S1.**
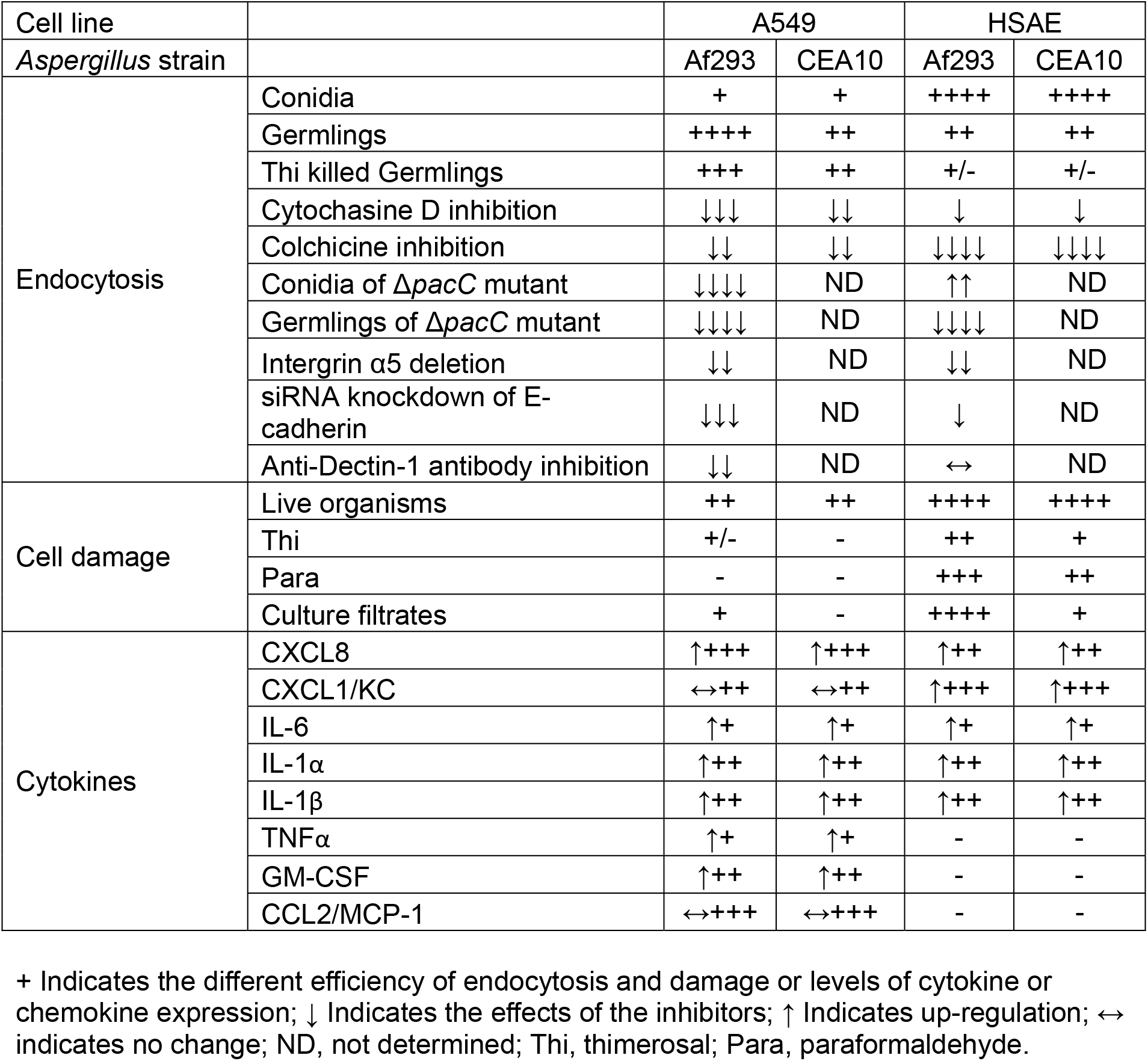
Summary of host cell endocytosis, damage, and cytokines or chemokines expression during A. fumigatus infection.

**Table S2.** A. fumigatus strains used in this study.

| Strain Name | Background | Genotype | Source |
| --- | --- | --- | --- |
| Af293 | Wild-type | Wild-type | P. Magee, University of Minnesota |
| CEA10 | Wild-type | Wild-type | N. Keller, University of Wisconsin-Madison |
| Af293-*GFP* | Af293 | *P<sub>gpdA</sub>-GFP-ble* | Liu and Lee et al., 2016 |
| CEA10-*GFP* | CEA10 | *P<sub>gpdA</sub>-GFP-ble* | Liu and Lee et al., 2016 |
| Δ*pacC* | Af293 | Δ*pacC::hph* | Present study |
| Δ*pacC-Comp* | Af293 | Δ*pacC::hph+pacC-ble* | Present study |
| Δ*pacC-GFP* | Δ*pacC*-Af293 | *P<sub>gpdA</sub>-GFP-ble* | Present study |
| Δ*pacC-Comp-GFP* | Δ*pacC-Comp*-Af293 | *P<sub>gpdA</sub>-GFP-ptrA* | Present study |

**Table S3.**
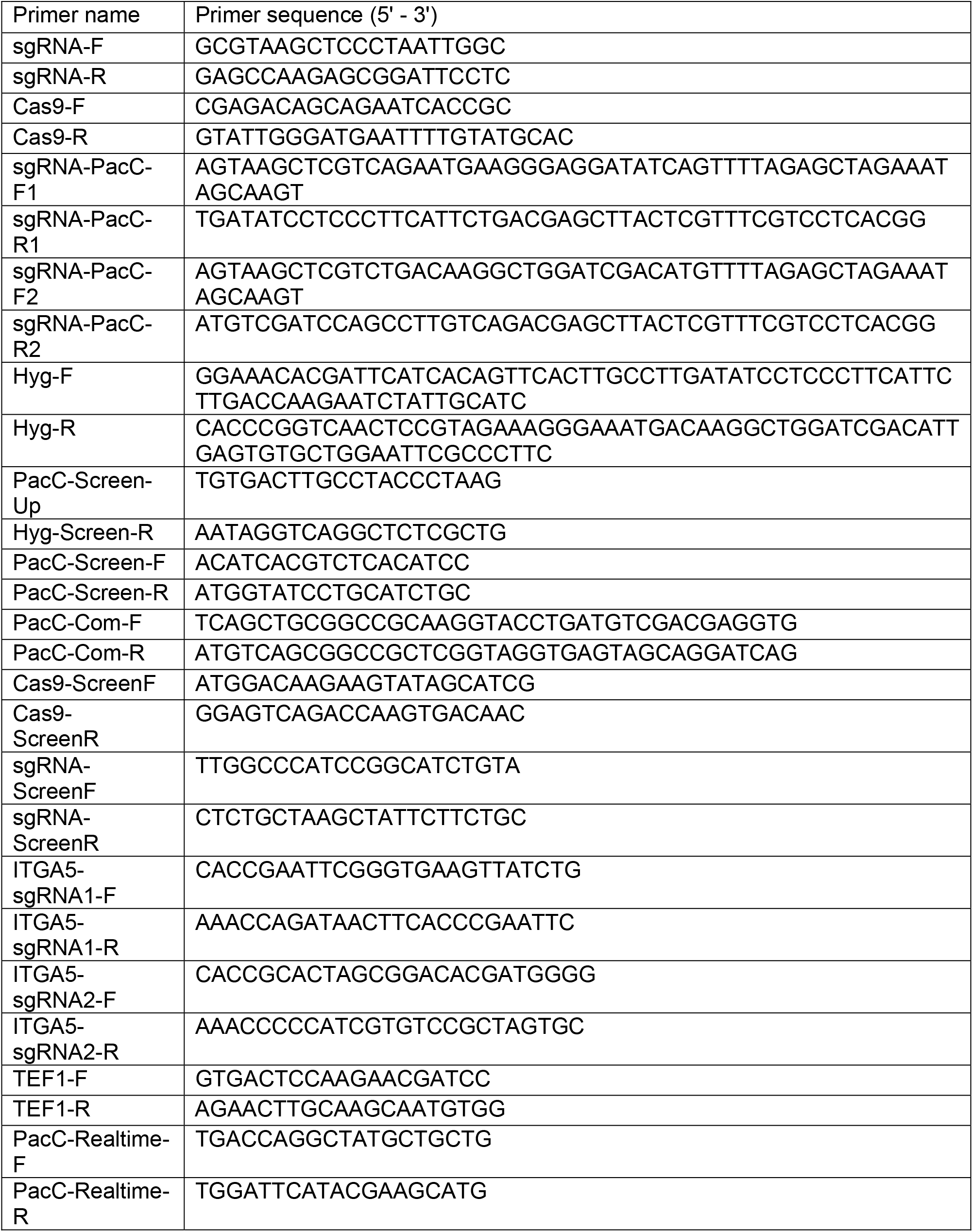
Oligonucleotides primers used in this study.

